# Sensitivity of estimates of the effectiveness of REDD+ projects to matching specifications and moving from pixels to polygons as the unit of analysis

**DOI:** 10.1101/2024.05.22.595326

**Authors:** Alejandro Guizar-Coutiño, David Coomes, Tom Swinfield, Julia P G Jones

## Abstract

There is a substantial interest in the potential of carbon credits generated by Reducing Emissions from tropical Deforestation and Degradation (REDD+) and traded on the voluntary carbon market for generating the finance needed to slow forest loss. However, such credits have become marred in controversy. Recent global-scale analysis using a range of methods for estimating the counterfactual rate of deforestation *ex post* suggest that many REDD+ projects have overestimated their effectiveness at reducing deforestation and consequently issued more credits than can be justified. All such methods include potentially arbitrary choices which can affect the estimate of the treatment effect. In addition, using pixels as the sampling unit, as some of the studies do, can introduce biases. One study which has been widely cited in the debate (Guizar-Coutiño et al. 2022) estimated avoided deforestation using statistical matching of pixels and a single set of matching options. We estimate avoided deforestation from the same set of projects using 7-hectare plots rather than pixels to sample deforestation and explore the sensitivity of the results to matching choices (exploring 120 matched sets in total). We filtered the results on three criteria: 1) post-matching covariate balance, 2) proportion of REDD+ samples that were successfully matched, and 3) similarity of trends in deforestation rates prior to REDD+ implementation (parallel trends). While one of the 44 REDD+ projects failed these quality control process, we estimate treatment effects for the remaining 43 projects. There was a substantial correlation between our new estimates and those published in Guizar-Coutiño et al. 2022 (0.72 measured in annual percent change, and 0.9 measured in total area change) and our headline estimate of 0.22% per yr (95% CI: 0.13 to 0.34) is essentially unchanged. At a time when confidence in the voluntary carbon markets is low, we hope these results provide reassurance that ex-post counterfactual estimates of avoided deforestation are consistent, helping accelerate their widespread adoption and rebuild trust in nature-based climate solutions.

## Introduction

Forests sequester and store enormous amounts of carbon, making forest conservation an important nature-based solution to climate change ^1^. REDD+ credits (Reducing Emissions from Deforestation and forest Degradation) are the most numerous of credits traded in the voluntary market ^2^ and there is very substantial interest in the potential of such credits to provide much-needed finance for tropical forest conservation ^3^. However, the extent to which REDD+ projects have provided the reductions in deforestation or forest degradation they have claimed has become hotly debated ^4–7^.

To sell carbon credits on the voluntary carbon market, a REDD+ project needs to demonstrate additionality: i.e. that any reduction in deforestation is due to the project ^8^. This requires an estimate of the counterfactual: how much deforestation would have occurred without the project ^4^. REDD+ projects do this by using a variety of methodologies, the majority published and administered by the nonprofit organization VERRA, to estimate ‘baseline’ deforestation. Baselines are based on historical deforestation in project reference areas which are selected to be as similar as possible to project areas in terms of drivers of deforestation ^9^. Deforestation in project areas is then compared with the baseline and, following verification, VCUs (voluntary carbon units) are issued which can then be traded on the voluntary market. This approach has advantages as allows a project to have some certainty up-front as to the number of credits a project can produce, however it has been criticized for putting a lot of weight on the ex-ante baseline estimates, rather than using information on real rates of deforestation in comparable areas during the project period to construct counterfactuals ^9,10^.

A suite of independent global-scale analyses using a variety of approaches have suggested that many REDD+ projects provide limited additional avoided deforestation. For example, Guizar-Coutiño et al. ^11^ found that 33 out of 40 projects studied did not avoid any deforestation. Even more concerning, there have been claims that projects have issued more voluntary carbon units than can be justified^10,12^. For example West et al.^10^, found that across 18 REDD+ projects for which they had data, projects had been used to offset almost three times more carbon emissions than their contribution to climate change mitigation, though these estimates are disputed ^7^.

While many analyses of the impact of conservation interventions on deforestation uses pixels as the unit of analysis (e.g. ^13–15)^ there is increasing recognition that analyzing deforestation as binary, unrepeatable data (a pixel is deforested or not) can result in biases which are reduced by aggregating binary pixels spatially ^16^. Where appropriate polygons exist, the unit of analysis can be pixels aggregated within land holdings ^17^, or administrative boundaries ^18,19^. However in many contexts appropriate units may not exist, or polygons reflecting such units may not be available. In such cases polygons of a given area placed at random can be used to sample the landscape ^10^

Given that forest conservation projects, including REDD+, are never randomly placed in a landscape, evaluating their impact on deforestation requires causal inference from observational data ^20^. Areas where REDD+ projects are set up (treatment units) will inevitably differ in terms of confounders which influence both exposure to the intervention (in this case the location of REDD+ projects) and the outcome of interest (deforestation). Statistical matching is often used to ensure treatment and selected control units are as similar as possible with respect to these observable confounders ^21,22^. However, there are a range of somewhat arbitrary modelling choices made when applying statistical matching which can influence the estimated treatment effect ^23,24^.

In this paper we revisit the same set of REDD+ projects as were presented in Guizar-Coutiño et al. ^11^. While Guizar-Coutiño et al. ^11^ sample deforestation using pixels, we instead sample deforestation using circular polygons of 150 m radius, or 7.1-ha plots. We also explore how the choices made during statistical matching (applying 120 different combinations of matching options to each project) influence the estimated treatment effect.

## Methods

The research pipeline is shown in Fig. 1 and developed in the following sections. Our outcome of interest is deforestation as reported in the Tropical Moist Forest (TMF) maps which present deforestation and degradation across the humid tropics at 30 m resolution from 1990 to 2019 ^25^.

**Figure 1.**
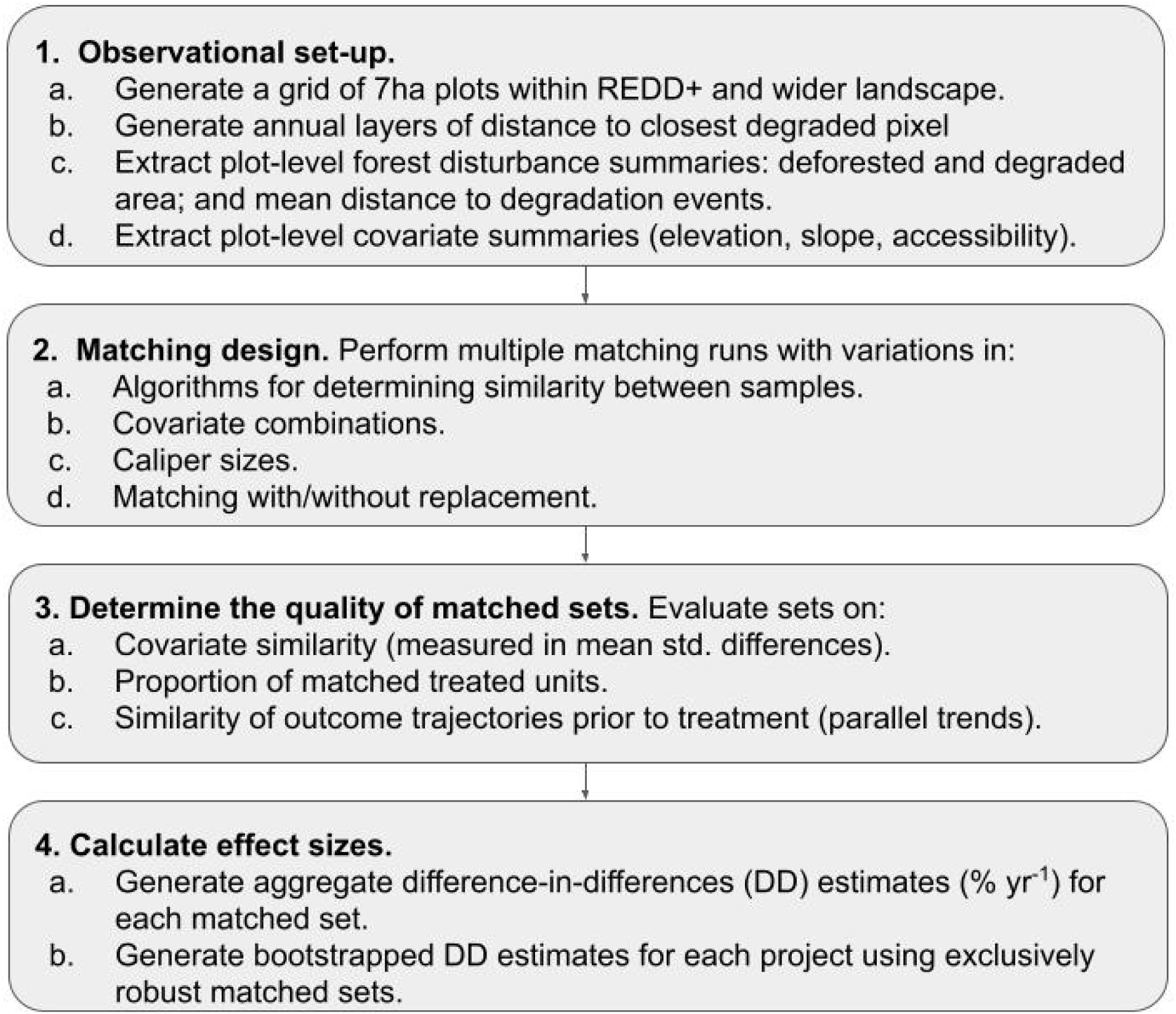
Schematic of the pipeline use to explore the impact of matching decisions on estimates of REDD+ effectiveness. Blocks indicate components of the analysis and are described in more detail in the methods.

### 1. Observational set-up

#### Selection of REDD+ projects

The VCS database is the leading accreditation registry for voluntary REDD+ projects ^2^ and the largest sources of data on voluntary REDD+ projects locations. We used the REDD+ database compiled by Guizar-Coutiño et al. ^11^, which identified 81 “Reducing Deforestation and Degradation” projects established in the tropics ^26^ and successfully found boundary data for 71 of these. While new projects have been introduced since 2019, and others withdrawn, for consistency with ^11^ we analyze the same set of projects. We excluded projects with <80% evergreen forest cover at the start (because the TMF dataset only covers evergreen forest), and projects which operated for fewer than 5 years or commenced before 2000. In Guizar-Coutiño et al. ^11^ we included project PL1748 in the analyses despite not meeting the minimum number of years for our criteria. This is because we recorded its start date incorrectly during our original analysis. We include it here for consistency despite it only running for four years.

#### Sampling within REDD+ areas and forested landscapes

To assess the extent of reductions in deforestation due to REDD+, we compared changes in forest cover within treated areas with those taking place in similar forest patches within the wider landscape. We generated plots with a radius of 150 m on a 1 km grid within forested areas of the REDD+ projects and surrounding regions. Each plot was 7.1 hectares in size and contained up to 78 TMF pixels, which we considered adequate for characterizing local drivers of deforestation and our outcome as continuous variables ^27^. We used the extent of forest cover in 1990 as delineated by the TMF ^25^ to position sampling plots, ensuring that all samples contained at least some portion of undisturbed moist forest cover when building the time-series of forest cover. We used the same approach to sample tropical moist forests plots in the wider landscape: we removed sampling locations that were not forested or that lay within a 15-km buffer zone around the project area, which may have been affected by local leakage of economic activities. We then random subsampled, until we had up to five samples in the wider landscape for each sampling plot located inside the REDD+ project, each separated by at least 1 km to reduce the spatial dependence ^28^; this created a pool of samples which was subsequently used to find matches for REDD+ project samples (Fig. 2).

**Figure 2.**
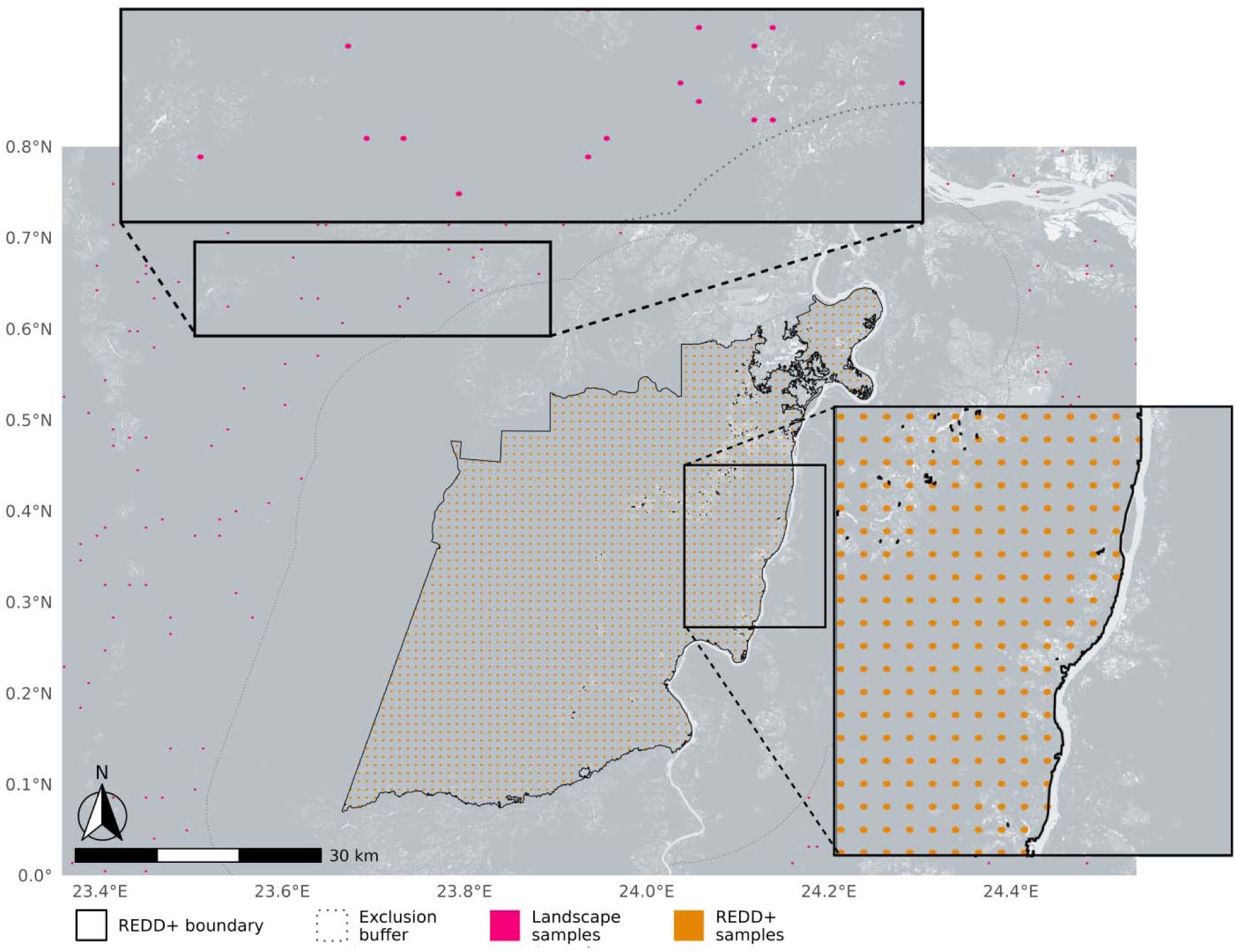
Illustration of the sampling strategy, for a REDD+ project in Democratic Republic of the Congo (VCS 1359). Features shown include the boundary of the REDD+ area (solid boundary) and the 7.1-ha plots collected from treatment (orange) and landscape (pink) areas. A buffer surrounding the project boundaries (dotted boundary) is shown within which we did not potential controls. The extent of undisturbed forest cover (dark grey) was used as a reference for distributing sample plots.

#### Plot-level forest disturbance estimates

Annual estimates of forest cover and forest disturbances were extracted from the TMF database. This database provides a long-term characterization of forest disturbances and land cover change, on an annual basis spanning the period 1990 -2019. Our analyses focused on examining temporal patterns of three main forest classes: The undisturbed class, which refer to closed evergreen or semi-evergreen forest areas that have not been degraded or deforested; the degraded class, characterized as short-term disturbance (less than 2.5 years); and the deforested class, representing a long-term removal of forest cover^25^. For each year during the 1990 - 2019 period, we calculated the extent of undisturbed, and disturbed forests classes within sampling plots. From these metrics we derived our outcome of interest - the proportion of forest lost - defined as, where Δp_t_ is the total area deforested in year t and p_d_ is the area of undisturbed forest in the baseline or reference year (see 4. Exploring the impact of matching choices on estimated treatment effect)

#### Drivers of deforestation

We identified key covariates associated with deforestation ^29^, some of which were also likely to have been associated with assignment to the treatment ^30,31^. To characterize deforestation risks factors, we collected data on key and observable bioclimatic and socio-demographic determinants of deforestation (as in ^29^ Table S1). For each circular plot, we obtained mean estimates for elevation and slope ^32^, distance to the nearest urban center ^33^and distance to forest edge. To account for distance to forest edge, we produced annual time-series of the distance to the nearest deforested pixel, as characterized by the TMF time-series. First, for each pixel covered by the circular plot, we computed pixel-level estimates of the distance to the nearest pixel that had changed its status from undisturbed to deforested, or from degraded to deforested, during the observed year, for the period 1990-2019. We then produced plot-level annual rolling estimates of the mean distance to deforestation events in the previous five years, covering the period 1995-2019.

### 2. Matching design

We used statistical matching to select a subset of samples in the wider landscape that were as similar as possible to samples in the project areas in terms of observable confounders: initial forest cover density, exposure to drivers of deforestation and trajectories of deforestation in the period prior to REDD+ (Table S1). We exact matched on country. We used a “base” set of covariates for all runs, which included: 1) elevation ^32^, 2) slope ^32^, 3) distance to population centers ^33^, 4) distance to degraded areas over the five years prior to project commencement, and 5) extent of degraded area at the time of project implementation.

To evaluate the sensitivity of estimates of avoided deforestation to matching specifications ^24^, we ran matching runs with alternative specifications of:

**Algorithms:** We used propensity score (PSM), Mahalanobis distance (MHN) and random forest (RFM) algorithms;

**Covariates:** We first used the “base” set of covariates alone, then iteratively added the following covariates to that base set: extent of undisturbed forest in 1990 (forest area 1990), extent of undisturbed forest in the project implementation year (forest area), and extent of undisturbed forest in 1990 and implementation year (forest area + forest area 1990); resulting in 4 distinct covariate combinations.

**Calipers:** We applied calipers of different sizes to constrain the selection of landscape observations prior to matching, so that the covariate distribution lay within 0.1, 0.3, 0.5, 0.7 and 0.9 SD of the covariate distribution in the treatment group.

**Matching with and without replacement:** Matching iterations allowed for both, matching with replacement (e.g. allowing control observations to be paired with multiple treatment units), or without replacement (e.g. constraining one control observation per treatment unit).

The combination of matching criteria resulted in 120 matching runs for each of the 44 selected REDD+ sites, giving a total of 5280 runs.

### 3. Assessing the quality of matched sets

Matching runs were assessed against three criteria:

**Covariate balance between treated and control samples.** Similarity between treated and control samples was determined by the overlap of the distributions of the covariate values. Matching runs were considered adequate if the absolute standardized mean difference between treated and matched control samples was less than 0.25 across all covariates included in the matching trial^22^.

**Proportion of project-area samples that were matched.** Estimates of additionality can be biased unless a high proportion of treatment samples are successfully matched: only then is the “average treatment effect on the treated” estimated accurately^22^. Matching was considered adequate if at least 80% of treatment samples were paired with a control.

**Parallel trends**. A key condition of difference-in-differences designs is that inferences cannot be drawn unless treatment and control groups follow similar outcome trajectories in the period prior to the intervention; this is known as the parallel trends condition ^34^. Since our sampling strategy involves characterizing historical deforestation within the sampling plots, counterfactuals would be credible only if deforestation trends between treated and control plots prior to the introduction of REDD+ are comparable. We established the existence of parallel trends by ruling out significant differences in rates of forest loss prior to project implementation (exemplified in Fig. S1). To assess parallel trends, a linear model was fitted for each project:

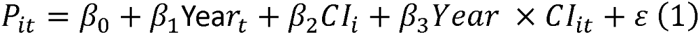

Where *P_it_* is the proportion of undisturbed forest area in the plot *i* in year *t*, and CI is the treatment group indicator (*i* =1 for treated or 0 for control), relative to forest extent in 1990. Parameter {3_3_ allows us to test for significant differences in deforestation trends between treated and control plots. We performed these tests using all available years before the project starting date. Departure from the parallel trends was assessed by testing whether the slope of the treatment and control groups were significantly different at p< 0.05; and matching was considered adequate if no departure was detected.

## 4. Calculating effect sizes

### Matched model effect sizes

To estimate additionality for each matched set, we calculated annual forest loss rates by aggregating the extent of deforested area across all sampling plots for each group (e.g. treatment or control) in the five years before and after project implementation, divided by the extent of undisturbed forest across all sampling plots in each group in the 6^th^ year prior to project implementation (i.e., reference year). We then estimated a mean difference-in-differences in forest loss rates as: DD = (AT + AC) – (BT + BC), where DD is difference-in-differences, A and B refer to the before and after period and T and C refer to the treatment and control group rates of deforestation rates (% yr^-1^).

### Bootstrapped effect sizes

We computed project-level mean difference-in-difference with 95% confidence intervals by bootstrapping difference-in-difference estimates of robustly matched sets. To produce a global estimate of the impact of REDD+ in reducing deforestation, we computed the mean of the site-level averages and estimated 95% confidence intervals by bootstrapping on the site-level averages.

## Results

### Sites included in the analysis

Of the 71 mapped REDD+ projects, 22 had less than 80% evergreen forest cover and 5 did not meet our time frame criteria. We applied matching to the remaining 44 projects (Fig. 3). Only 22% of all matched sets met all three of our quality-control criteria (1142 of 5280 matched sets). One of the 44 REDD+ projects failed to produce any high-quality matched set (project 1201). The remaining 43 REDD+ sites were located in America (n=34), Africa (n=6) and Asia-Pacific (n=3).

**Figure 3.**
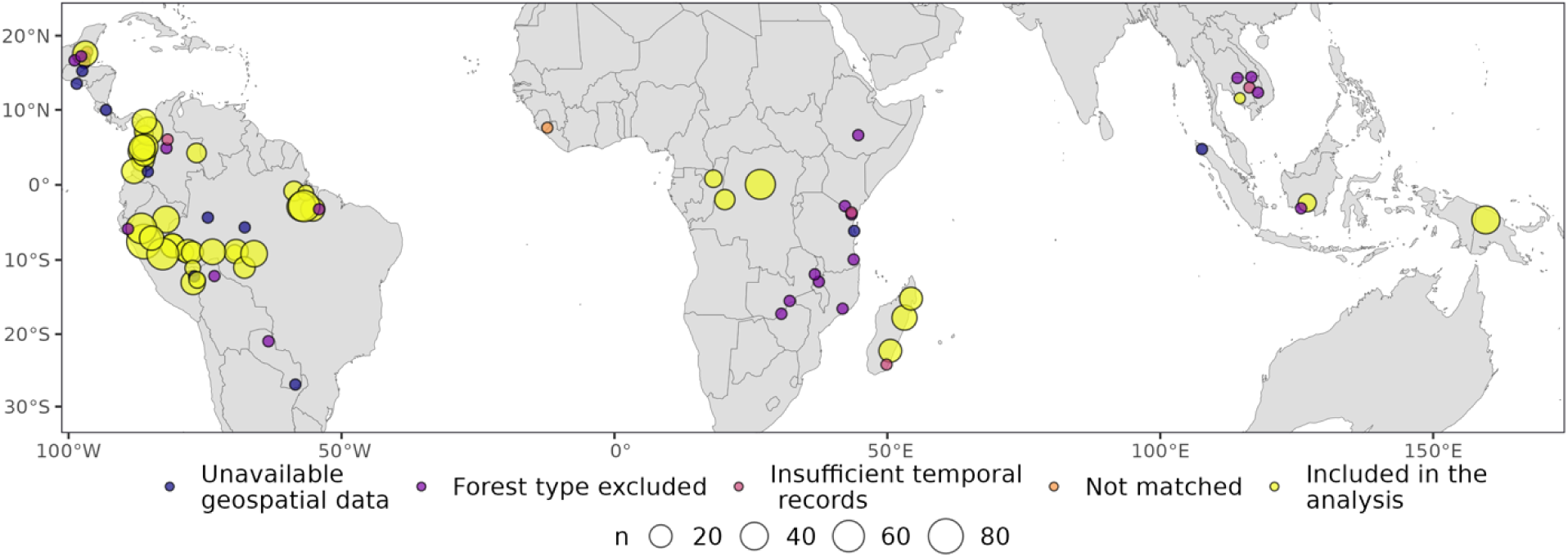
Location of the REDD+ projects included in the analysis. Of the 71 mapped sites, 22 had less than 80% evergreen forest cover at the start of projects (purple), 5 did not meet our time frame criteria, leaving 44 projects that were examined with multiple matching approaches (34 in the Americas, 7 in Africa and 3 in Asia-Pacific). Successful matching models were returned for 43 projects (yellow), with plots size indicate the number of robust models available for each of them. Project PL1748 did not met the time frame criteria but was included in the analyses for consistency with Guizar-Coutiño et al. 2022.

### Matching parameters and quality of matched sets

The quality of matched sets was more influenced by the choice of algorithms and by covariate selection, than by caliper size or matching with or without replacement. We note varying success rates across the three matching quality criteria (Fig. S2): 70% of sets failed the parallel trends criteria (i.e. they did not display similar outcome trajectories before the REDD+ project was established; Fig. S1), while 54% failed to achieve covariate balance between treated and control observations, and 31% failed to meet the criteria that 80% of treatment plots were successfully matched. Matched sets using the Mahalanobis distance method were much more likely to pass the three matching quality tests than sets matched with other methods (Fig. 4). Moreover, including measures of forest area on different time periods alongside the base covariates increased the quality of matched sets. Neither the caliper size nor the choice of whether to match with replacement had much effect on whether a matched set passed the quality control tests.

**Figure 4.**
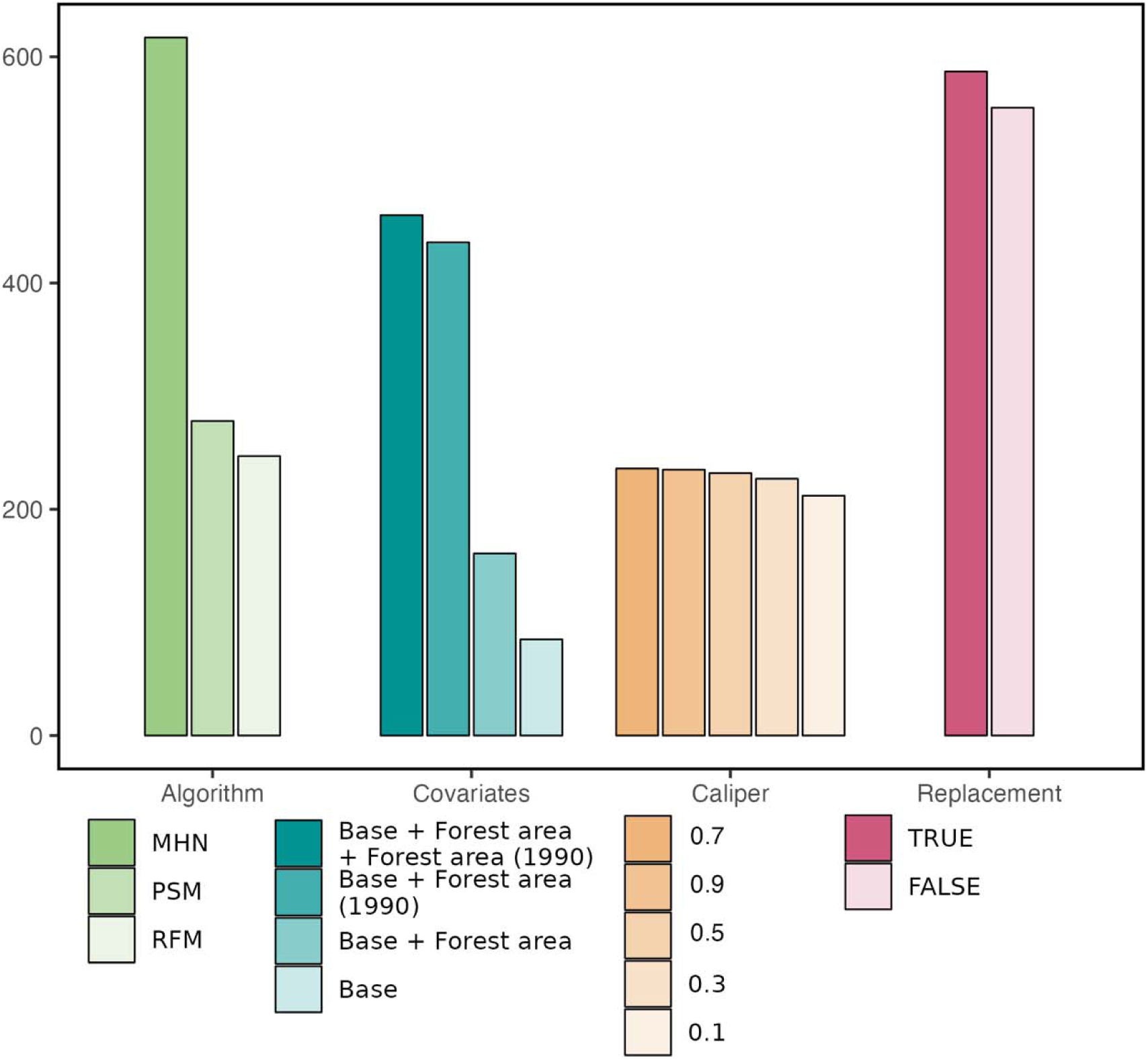
The number of matched sets that passed all three quality control criteria, subdivided by specifications made when matching. A total of 120 alternative matching specifications were made in each of 44 REDD+ sites, giving a total of 5280 matching sets; of these 1142 met all three quality control criteria. The specifications included: a) four choice of covariates combinations; base covariates only (i.e. elevation, slope, accessibility, distance to degradation, country and biome), base covariates plus forest cover at the time of project implementation, base covariates plus forest cover in 1990, and base covariates + forest cover in 1990 and implementation year; b) choice of matching algorithm between Mahalanobis (MHN), propensity score (PSM), and random forest (RFM); c) choice of filtering regime (caliper sizes 0.1, 0.3, 0.5, 0.7 and 0.9 SD); and d) choice of whether to allow matching with replacement or not.

### Revised estimates of effectiveness of REDD+ in reducing deforestation

Our analyses highlights the variability in the avoided deforestation delivered by REDD+ projects over their first five years of operation, both in terms of relative reductions in forest loss (Fig. 5a), and absolute deforestation avoided (Fig. 5b). While some projects have delivered substantial avoided deforestation (those to the left hand side of the Figure), the majority have effect sizes close to zero. Our project-level difference-in-differences estimates of robust matched sets were well correlated with the estimates produced by pixel-matching in Guizar-Coutiño et al. ^11^. The correlation is 0.72 when measured in annual percent change, and 0.9 when measured in total area change.

**Figure 5.**
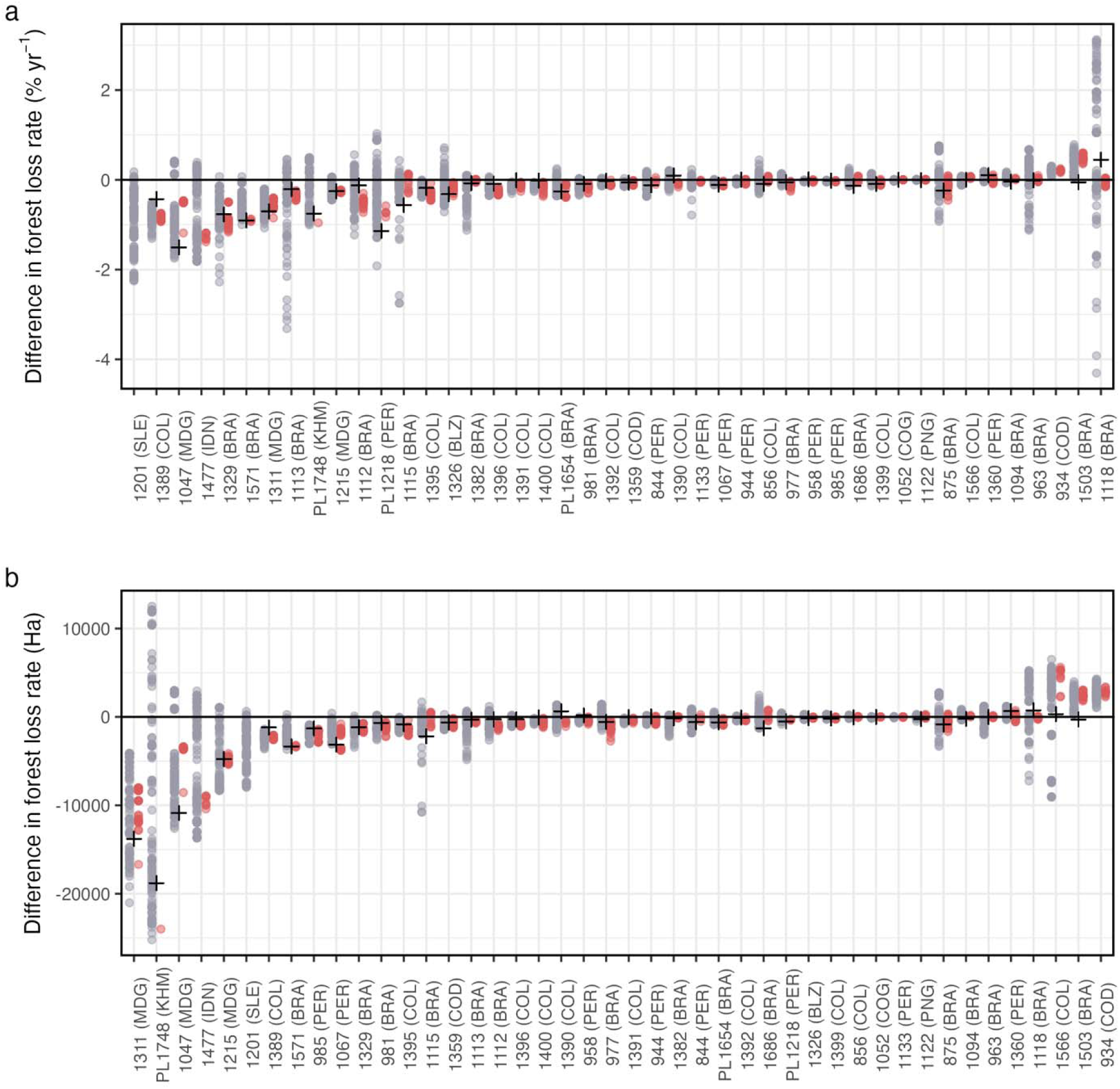
Annual differences in (a) deforestation rates for the 44 REDD+ projects included in this study and (b) total area loss using all matched models, with estimates below the zero-line indicating avoided deforestation. Dots indicate difference-in-differences estimates coloured according to whether the matching met our quality control criteria (red=robustly matched, grey=rejected). There are no valid models for project 1201 in Sierra Leone. For projects included in ^11^, estimates from that analysis are indicated with a ‘+’ symbol.

Table S2 presents the bootstrapped estimates of the deforestation avoided by each project (both in terms of percentage per year and hectares per year) with 95% confidence intervals. For 30 of the 43 projects, our best estimate of avoided deforestation as a percentage and total hectares is negative and does not overlap zero (Table S2). However, for many of those projects the reduction in deforestation is modest.

Combining the 43 site-level estimates, we observe a mean reduction of 0.22% yr^-1^ (95% CI: 0.13–0.36) in the rates of forest loss compared to controls, which amounted to an approximate area of avoided deforestation of 64903 Ha. This is essentially unchanged from the figures presented in Guizar-Coutiño et al. ^11^, which similarly calculated a mean reduction of 0.22%/year (95% CI: 0.13–0.34) amounting to approximately 66754 ha of avoided deforestation.

## Discussion

Given heated debate about the extent to which REDD+ projects have effectively avoided deforestation ^4,7,10,12,35^, the inevitable uncertainty in estimates using a single matching approach ^22,24^, and concerns about analyzing deforestation as a binary process ^16^, revisiting and updating the estimates of avoided deforestation provided in Guizar-Coutiño et al. ^11^ is valuable. We find that most projects reduce deforestation. However, effectiveness is highly variable; for many projects the reductions in deforestation are modest and for some they are non-existent. These revised estimates add to the growing evidence base exploring the impacts of site-based REDD+ project effectiveness using ex post counterfactuals ^10–12,36^.

Our previous study ^11^ used ∼30m pixels to sample deforestation. Our new approach, using 7.1-ha plots all separated by at least 1 km, has the advantage of reducing the spatial dependence of samples ^28^. By using plots as our sampling unit (within which deforestation is a continuous variable) rather than pixels (where deforestation is a binary outcome), we were also able to select controls with similar trends in deforestation. There are inevitably a range of somewhat arbitrary choices which need to be made when selecting matching approaches ^21,22^. By trialing 120 combinations of matching options and estimating the treatment effect from the full set which meet quality-control criteria, we get a valuable estimate of the uncertainty in estimates of ex post estimates of avoided deforestation.

This work explores sensitivity to how matching on observed confounders was carried out. However, there is a risk that important confounders remain which cannot be accounted for ^37^. Despite the importance of testing for sensitivity of results to unobserved confounding, this is not yet standard practice in conservation impact evaluation ^38^. In the context of REDD+, hidden confounders might include institutional factors such as tenure security and presence of local conservation initiatives ^39–41^ or local drivers of agricultural activities ^29^. Some of the critiques of the recent published estimates of the avoided deforestation from REDD+ projects ^7,42,43^ suggest that such large-scale analysis (covering multiple REDD+ projects in multiple sites using the same methodology) inevitably fails to account for key local-level drivers of deforestation. Further work exploring the sensitivity of these results to hidden confounders would be valuable.

Climate change is advancing faster than many climate scientists had predicted. The target of keeping warming within 1.5 degrees of pre-industrial levels has already been missed ^44^, or will be soon ^45^. Given the contribution of tropical forests to both sequestering and storing carbon^46,47^, it is vital that the global community find ways to stop further loss of tropical forests ^4^. The voluntary carbon market is not the only way that forest conservation can be funded. International policy is focused on developing a UN-governed schemed under article 6 of the Paris Climate Agreement which will focus on lowering deforestation over entire jurisdictions ^40^ rather than in specific REDD+ project locations. There are also many bilateral agreements where countries support other countries to lower their deforestation-related emissions.

However, selling emissions reductions from avoiding deforestation on the voluntary carbon market is generating substantial, and hard-to-replace, finance for forest conservation and will remain important at least into the medium term. The lack of credibility of the credits being issued poses a threat to this important revenue stream ^35,48^. There is growing recognition that conservative credits issuance ^49^ can help address issues such as impermanence ^50^, and leakage ^51^. At time when confidence in the voluntary carbon markets is low, we hope our results provide reassurance that *ex-post* counterfactual estimates of avoided deforestation can deliver relatively consistent estimates of additionality, helping accelerate their widespread adoption and rebuild trust in nature-based climate solutions.

## Acknowledgments

A.G.C. received a scholarship from Consejo Nacional de Ciencia y Tecnología México, CONACYT, and the Frank Jackson Foundation through a grant to Wolfson College, as well as funding from the Tezos Foundation gift to the Cambridge Centre for Carbon Credits (4C). J.P.G.J. was an Oxford Martin School visiting fellow and short-term visiting fellow at Jesus College Oxford when working on this paper.

## Data and code

Data summaries and code to reproduce the main figures provided at: https://github.com/guizar/redd-sens

## SUPPLEMENTARY INFORMATION

### SUPPLEMENTARY FIGURES

**Figure S1.**
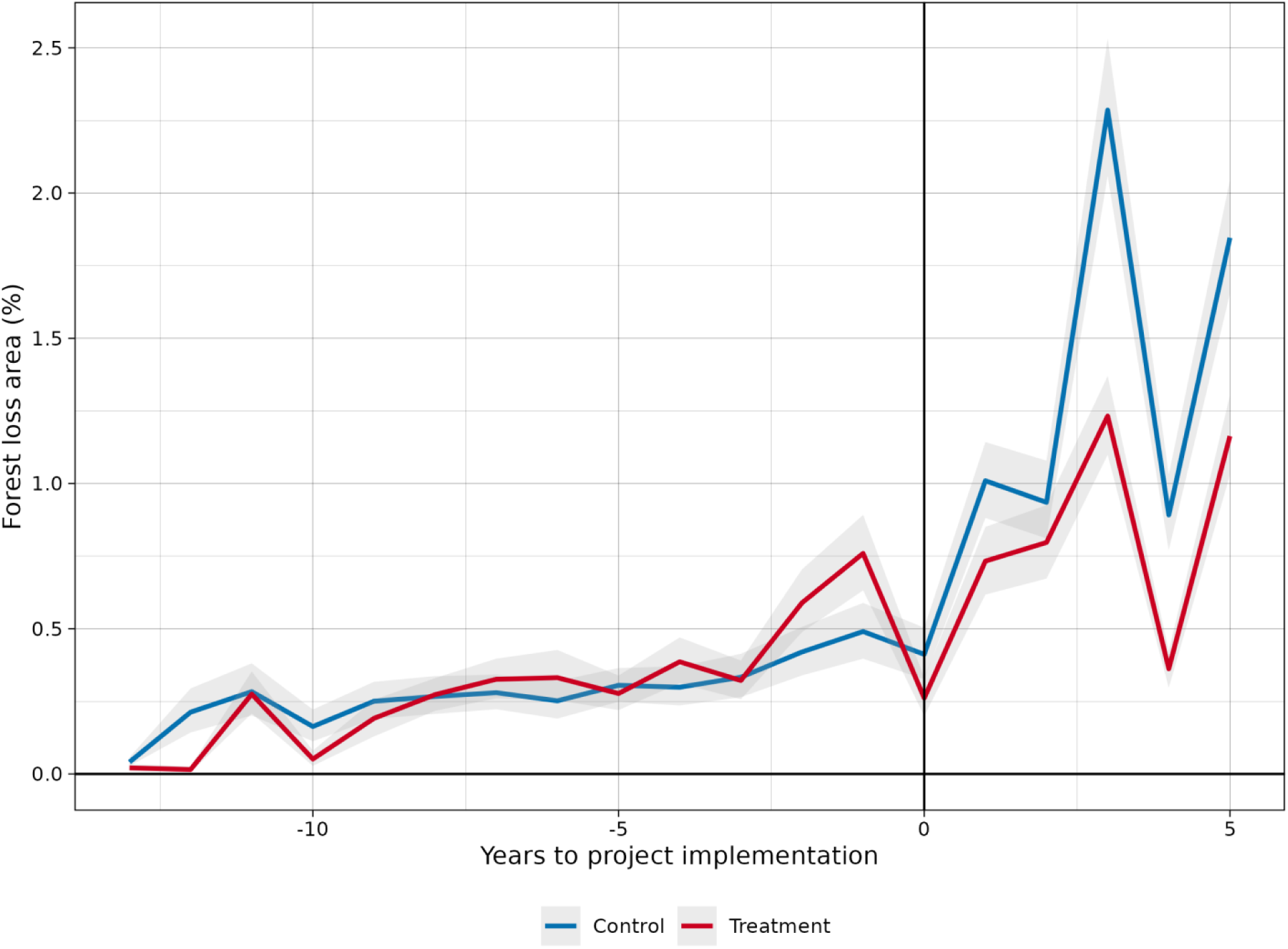
Patterns of deforestation, before and after project implementation, for the Ankeniheny-Zahamena (CAZ) Protected Area project in Madagascar (VCS 1311). Plot-level mean estimates of forest loss (Ha) in treated (blue) and control (red) samples. 95% confidence intervals (grey) generated by bootstrapping. This particular time-series comes from a matched set that meets all quality criteria and was used to assess the sensitivity of estimates to unobserved confounders.

**Figure S2.**
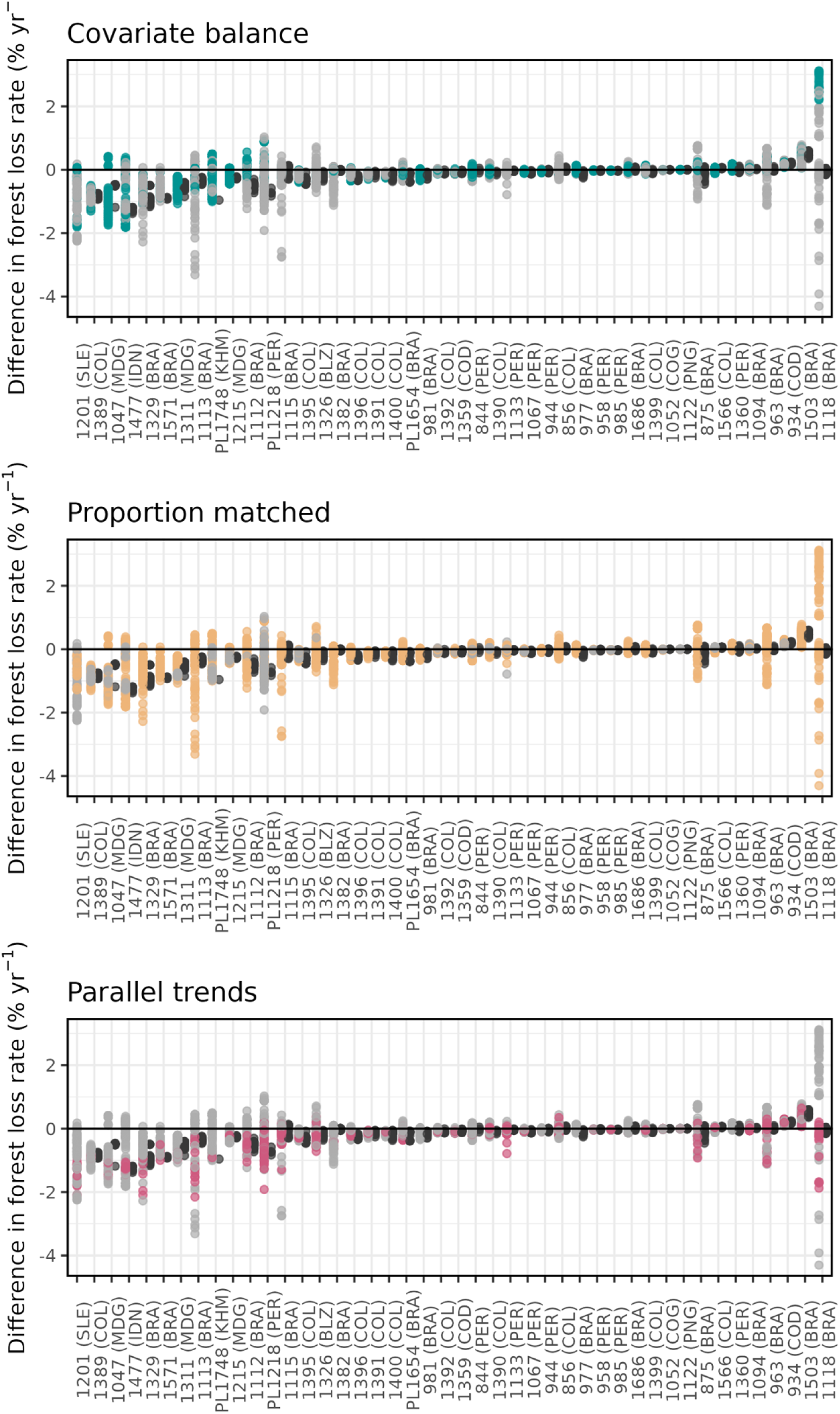
The relative importance of the three validation criteria as the reasons matched sets were rejected. The three panels show differences in deforestation rates for the 44 REDD+ projects (% yr^-1^) during the first five years of project implementation, arranged by project-level average differences (x-axis). Grey indicates that the run failed to meet the criteria covered by the panel, colour shows that it met the criteria but failed to meet other criteria, black indicates that all criteria were met.

### SUPPLEMENTARY TABLES

**Table S1:** Covariates selected for matching. All the covariates listed in the table are included as part of our core set of matching variables, except for ’Temporal estimates of forest cover’ variables, which we 460 used to iterate in the matching runs (shown in bold).

| Category | Rationale for inclusion | Source | Variables | Format and resolution |
| --- | --- | --- | --- | --- |
| Physical characteristics | Elevation and slope influence land use practices, with higher elevation and steeper slopes consistently associated with a lower risk of deforestation (Joppa & Pfaff 2009). | SRTM Digital Elevation Data Version 4 | Elevation (meters)<br>Slope (degrees) | Raster (90m2) |
|  |  | (Jarvis et al. 2008) |  |  |
| Accessibility | Accessibility to forest lands from urban centres is associated with a higher risk of deforestation (Chomitz et al. 1999; Rudel et al. 2009). | Global accessibility map (Weiss et al. 2018) | Time travel to population centers (seconds) | Raster (300m2)<br>Temporal res: 2015 |
| Bioclimatic | The biophysical characteristics of biomes, such as temperature, precipitation and agricultural suitability (Busch & Ferretti-Gallon 2023). Our sampling occurred in tropical moist forest regions for climatic comparability | JRC Tropical Moist Forest database (Vancutsem et al. 2021) | Biome (Tropical moist forest) | Raster (30m2) |
| Governance / political economy | Deforestation is influenced by domestic policies, governance, economic processes and markets (Umemiya et al. 2010; Ceddia et al. 2014). | Large Scale International Boundary (LSIB) dataset | Country | Vector |
| Proximity to deforestation | Proximity to cleared lands is consistently associated with higher deforestation risk (Busch & Ferretti-Gallon 2023). We computed the average distance to the closest deforested pixel in sampled plots used for matching | Own analysis using the JRC Tropical Moist Forest database (Vancutsem et al. 2021) | Euclidean distance to deforested areas (meters) | Raster (30m2) |
| Are of forest disturbance | The risk of deforestation increases when there are cleared or disturbed patches of forest (Busch & Ferretti-Gallon 2023). We calculated the area of degraded forests within plots by the year the project was started. | Own analysis using the JRC Tropical Moist Forest database (Vancutsem et al. 2021) | Degraded area (Ha) | Raster (30m2) |
| Temporal estimates of forest cover | We used estimates of plot-level forest cover at the beginning of the TMF time series (1990) and at the time of project implementation to ensure temporal dynamics of deforestation were captured in the matching | JRC Tropical Moist Forest database (Vancutsem et al. 2021) | Undisturbed forest cover extent (Ha) | Raster (30m2) |

**Table S2:**
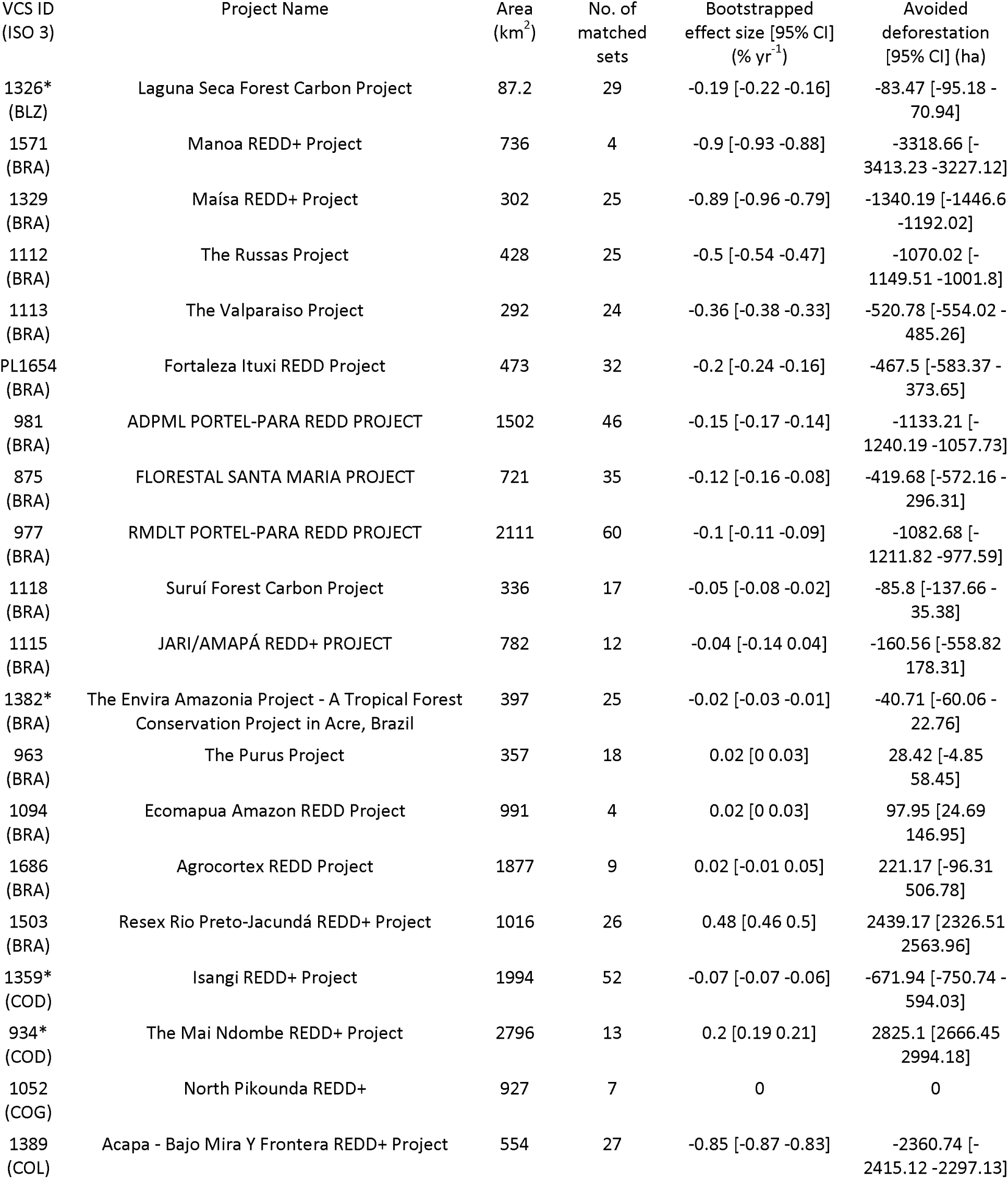
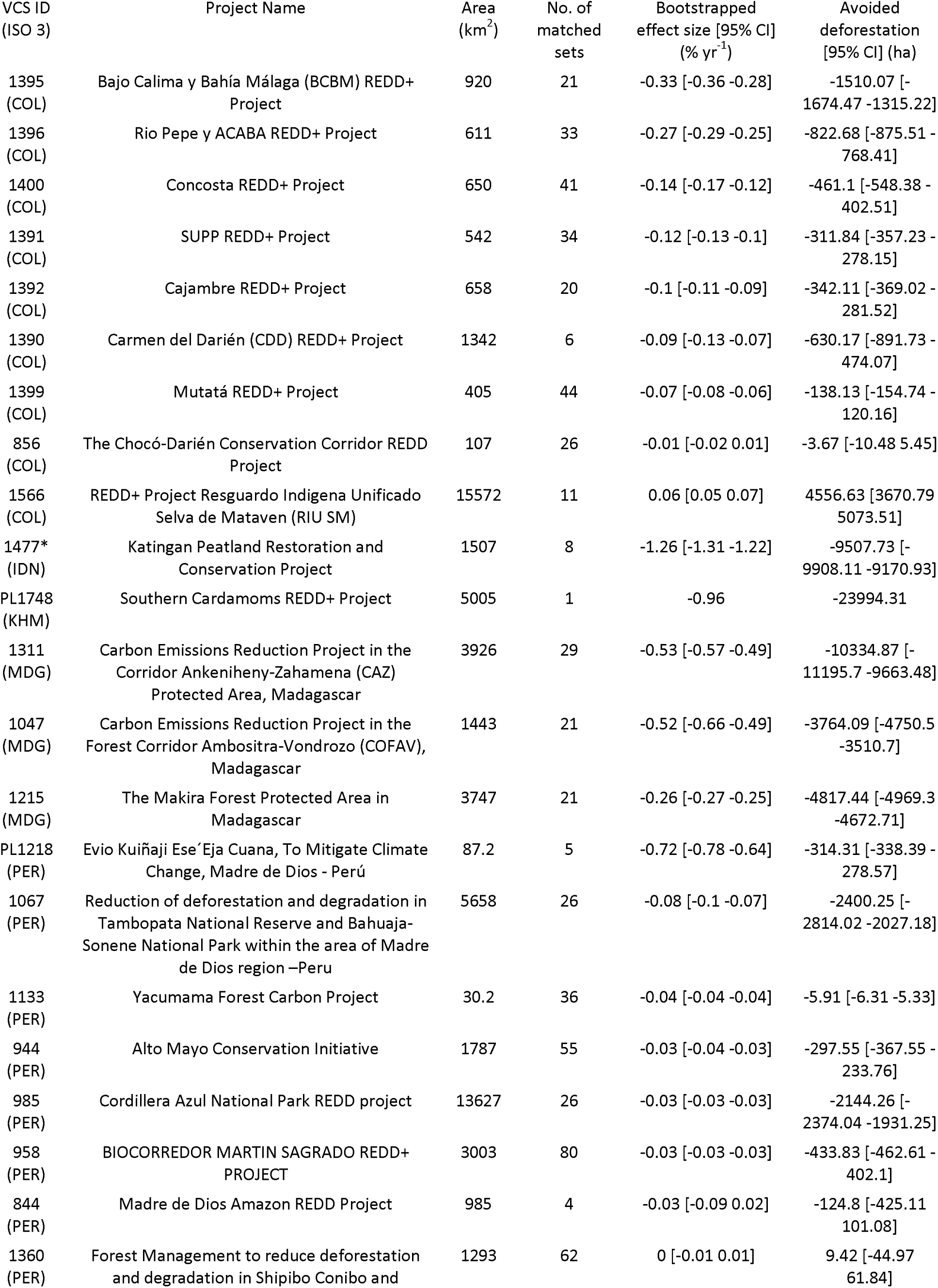
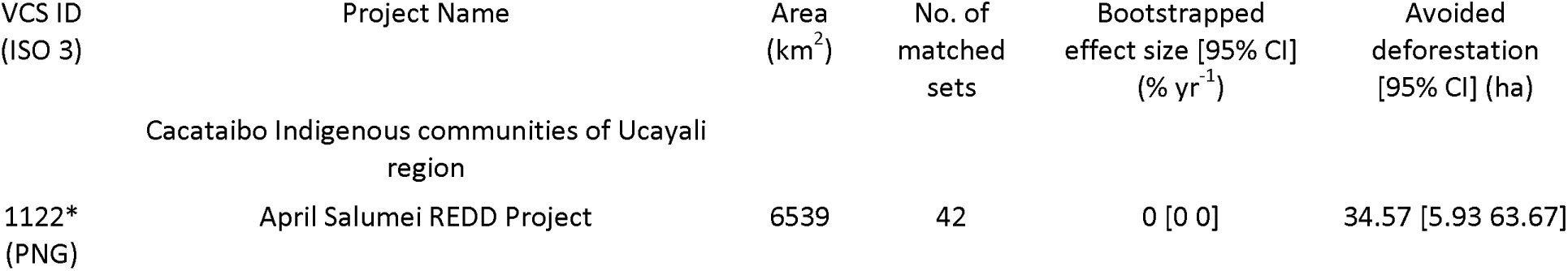
Project-level treatment effect estimates (bootstrapped robustly matched sets only) for the REDD+ projects for which valid matched sets are available. Estimated treated effects are averaged over the first 5 years. Projects classified as “Avoided Planned Deforestation and forest Degradation” indicated with asterisks.

